# The female larval fly CNS connectome: first draft

**DOI:** 10.64898/2026.09.28.754924

**Authors:** Samia Mohinta, Daniel Franco-Barranco, Shi Yan Lee, Pedro Gómez-Gálvez, Shaurya Agrawal, Michael Clayton, Christopher L. Barnes, Nadine Randel, Ana Correia, Nicolo Ceffa, Nadia Stoichkova Velichkova, Katrin Nielsen, Harald Hess, Song Pang, C. Shan Xu, Jimena Berni, Michał Januszewski, Marta Zlatic, Albert Cardona

## Abstract

We release an enhanced FIB-SEM volume and associated neuron, mitochondria, synapse and glia automated segmentations for the complete, gap- and artifact-free central nervous system of the first instar female larval *Drosophila melanogaster*. Segmentation data is largely unproofread besides select neurons in the premotor and proprioceptive system and select subvolumes where synapses have been manually annotated, serving as training and evaluation data. The dataset represents the first complete central nervous system of the larval fly, and now awaits substantial improvement of machine vision approaches that effectively leverage domain knowledge for the automated yet data-guided proofreading, or substantial and costly human proofreading. All the data and methods are publicly available for further improvement and analysis.

## 1 Introduction

Even genetically isogenic individuals present different synaptic wiring diagrams, or connectomes, as established in the nematode *C. elegans* with the analysis of the overlapping sections of the five different worms used to reconstruct its connectome [1]. In order to compare neural circuits across sexes or closely related species, in search of correlations between changes in circuit architecture with changes in circuit function or behaviour, we must therefore first measure the natural variability in synaptic connectivity, by mapping the connectome of multiple individuals. For the adult *Drosophila melanogaster* brain, three female brain hemispheres [2, 3] and two male ones [4] exist, with the variability having been analysed albeit in limited capacity given the very low number of samples [5].

Here, we release the complete reconstruction of the central nervous system (CNS) of the larval *Drosophila melanogaster*, largely unproofread, with its brain being the second fully mapped first instar female larval brain [6], adding up to a total of four brain hemispheres given the bilateral symmetry. The data, approach, and software tools reported here define a path forward for the mapping of the many more connectomes of both male and female larvae of *D. melanogaster* and other species of the genus *Drosophila* necessary to establish the natural variability of the larval CNS connectome, on the basis of which then identify and quantify the shared and the differing aspects of the connectome across individuals, sexes and species.

On the basis of the measured connectivity variability among identified neuronal cell types, a comparison across species could reveal which circuits are more readily evolvable, and establish how changes in circuit structure correspond to changes in circuit function. An example is the pioneering study comparing larval antennal lobes across *D. melanogaster* and *D. erecta* that identified very specific and functionally significant changes in the antennal lobe circuits of *D. erecta* [7] relative to *D. melanogaster* [8]. Another example is the identification of changes in the circuit for multisensory integration in the nerve cord of *D. santomea* [9] relative to *D. melanogaster* [10], alongside a functional intervention that demonstrates how the circuit of one species can be made to operate like that of the other. Yet a third example establishes how the directed genetic manipulation of synaptic input fractions in a multisensory integration neuron of the larval nerve cord leads to changes in the integration function of a neuron and therefore to changes in circuit function and behavioural responses to the same stimuli [11]. Collectively these studies illustrate the extraordinary power of the *Drosophila* model for experimental and comparative connectomics, with the reduced dimensions and reduced numerical complexity of the larva offering unique and specific advantages.

The volume of the whole female larval CNS, codenamed Octo, was prepared with an early iteration of the SuperFly sample preparation protocol [12] and imaged in 2018 with enhanced FIB-SEM [13] at 8×8×8 nm/pixel. The image volume is virtually free of artifacts, with no serial section loss, no curtaining artifacts, and no known membrane nicks that would result in artefactually merged neurons upon automated segmentation. The completeness of the data permits now the mapping and analysis of the complete female larval connectome, necessary to study how the brain controls the body via descending neurons, and how e.g., ascending proprioceptive feedback imposes constraints on behavioural choice by limiting the range of possible motor actions.

The volume is not proofread beyond a few select neurons in the premotor system and among ascending projection neurons of the proprioceptive system. Proofreading the complete volume will require either tens of man-years of effort or significantly improved superpixel agglomeration methods, yet to be developed by us or others. Such new methods could leverage existing knowledge of the spatial distribution of mitochondria to filter false-positive synapse detections given their known spatial correlation to presynaptic sites [14] and the relationship between mitochondrial morphology and neuronal cell types [15, 16]; the predictable and constrained extent and angle of microtubules, fully continuous from the neuron’s soma to throughout the axonal and dendritic arbors except for the most distal twigs (the fly’s dendritic spines; [14]) and readily recoverable from volume EM [17]; and the known spatial distribution of postsynaptic sites in microtubule-free distal twigs [18], with the median twig hosting only one postsynaptic site in first instar larval neurons [14]. On the meshes of the reconstructed larval neurons released here, new methods for analysing the distinct morphology of spines may prove profitable for automating various aspects of proofreading and cell type classification [19].

Alongside the EM volume, we release three different volumetric neuron instance segmentations with different spatial coverage, predicted pre–postsynaptic connections, mitochondrial instance segmentations, and a semantic segmentation of glial tissue. These data channels can be visualized together as multiple layers in Neuroglancer [20]. The whole-CNS neuron segmentation is additionally served through the Connectome Annotation Versioning Engine (CAVE), which supports collaborative proofreading and versioned refinement of large connectomic segmentations [21].

We evaluate the released reconstructions against Octo reference manual annotations and a modicum of manually-proofread neurons. Depending on the reconstruction layer, the corresponding ground truth was obtained through manual annotation, neuronal proofreading, or automated curation procedures, as described in the relevant sections below. To evaluate the accuracy, we quantitatively assessed long-range neuron continuity, synaptic partner detection on held-out Octo regions, mitochondria instance segmentation, and glia semantic segmentation, and provide qualitative examples of the resulting reconstructions. Together, the Octo eFIB-SEM volume and its associated reconstructions provide the first complete larval CNS volume and connectome, albeit pending full prooreading, and a second first instar female larval brain.

## 2 Volume Acquisition and Preprocessing

The Octo central nervous system (CNS) volume was acquired using enhanced focused ion beam scanning electron microscopy (eFIB-SEM) at 8 *×* 8 *×* 8 nm isotropic resolution. The final volume contains 20,486 *×* 21,471 *×* 34,745 voxels in *x, y, z*, corresponding to a physical extent of approximately 164 *×* 172 *×* 278 *µ*m.

Following acquisition, the image volume was registered using a custom Fiji/ImageJ workflow developed for the laboratory’s FIB-SEM data.^1^ The workflow incorporates the elastic registration framework of Saalfeld et al. [22]. Local correspondences between image planes are estimated by block matching and regularized using an elastic model to correct residual misalignment while limiting artificial deformation [22]. The resulting transformations were applied to assemble the registered three-dimensional Octo volume.

After registration, contrast-limited adaptive histogram equalization (CLAHE) was applied to enhance local image contrast [23]. This registered and CLAHE-processed volume was used for the reconstruction and analysis described in the following sections.

## 3 Automated Reconstruction Layers

In addition to the EM volume, we release automated neuron and mitochondria instance segmentations, a semantic segmentation of glial tissue, and predicted synaptic connections. The FFN neuron segmentation, synaptic predictions, mitochondria segmentation, and glia segmentation span the complete Octo CNS, while the LSD- and BiaPy-based neuron segmentations are restricted to the neuropil.

All of these outputs were generated using machine-learning models applied to the Octo EM volume. Below, we describe how each reconstruction was generated and evaluate its agreement with Octo reference annotations and proofread neurons.

### 3.1 Neuron Segmentation

Octo includes three neuron instance segmentations generated using Flood-Filling Networks (FFNs) [24], Local Shape Descriptors (LSDs) [25], and a BiaPy-based affinity model [26]. The FFN segmentation covers both neuropil and soma regions across the complete central nervous system (CNS), whereas the LSD- and BiaPy-based segmentations cover the neuropil throughout the CNS. This difference in coverage is illustrated in Fig. 1.

**Fig. 1:**
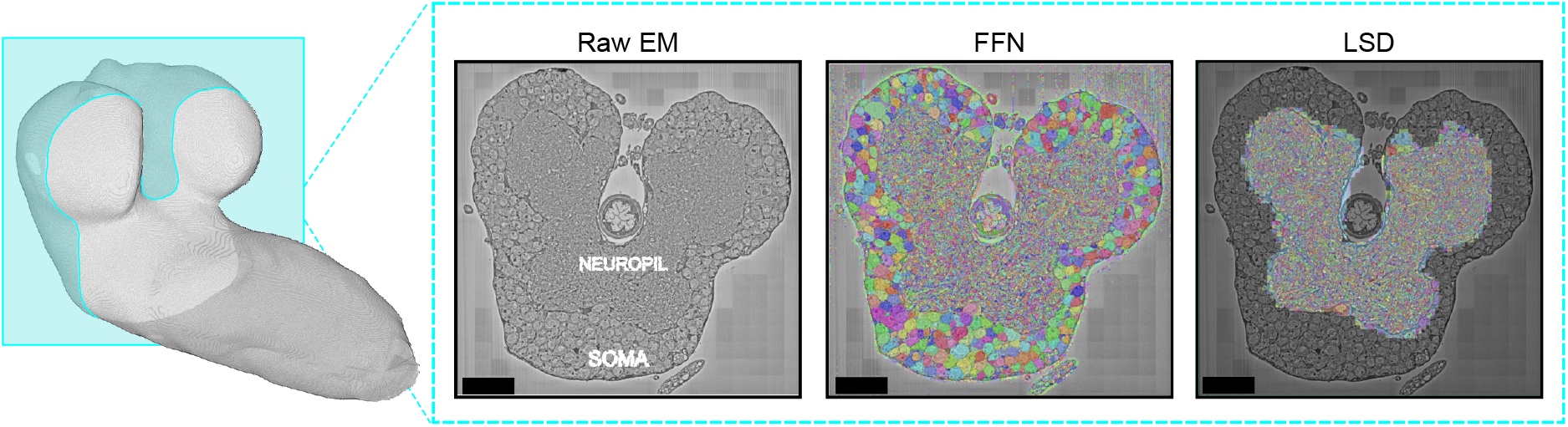
Spatial coverage of the neuron segmentations in Octo. Left, three-dimensional view of the Octo CNS with a cross-section shown in cyan. Right, the corresponding raw EM section, with neuropil and soma regions indicated, together with the FFN and LSD segmentations. The FFN segmentation covers both the neuropil and surrounding soma regions, whereas the LSD segmentation is restricted to the neuropil. Scale bar: 20 *µ*m.

#### Flood-Filling Network-based segmentation

The whole-CNS segmentation was generated in collaboration with the Google Connectomics team using FFNs [24].^2^ FFNs reconstruct neuronal instances by iteratively extending an object mask through overlapping local fields of view. The resulting segmentation covers both neuropil and soma regions across the complete CNS (Fig. 1). We use this segmentation to assign predicted synaptic sites to neuronal segments and obtain the current whole-CNS connectivity graph. The FFN segmentation also provides the starting point for whole-neuron proofreading.

#### Local Shape Descriptors-based segmentation

A second neuron segmentation was generated using Local Shape Descriptors (LSDs) [25]. We use the PyTorch implementation of LSDs released in Catena [27].^3^ The model jointly predicts local shape descriptors and voxel affinities. The predicted affinities were converted into neuron instances using watershed followed by WATERZ agglomeration [28]. Training used the publicly available Hemibrain ground-truth volumes released with the original LSD code by Sheridan et al. [2, 25]. Inference was run across the neuropil regions of the complete Octo CNS (Fig. 1).

#### BiaPy-based segmentation

The third neuron segmentation was generated using an affinity-prediction model implemented in BiaPy [26]. Training data were derived from the Octo FFN segmentation by extracting 356 three-dimensional crops distributed throughout the neuropil. The corresponding FFN labels were curated automatically before training. This processing removed small spurious segments, merged nested or fully enclosed fragments, filled internal holes, and corrected cases in which intracellular regions, including mitochondria, had been assigned separate instance IDs within a neuron. The resulting labels were used as pseudo-labels for model training.

A pretrained STU-Net [29], initialized using OrgMIM pretraining [30], was trained within BiaPy to predict voxel affinities from the Octo EM. The predicted affinities were converted into an initial segmentation using the watershed implementation in `skimage.segmentation` [31], followed by WATERZ agglomeration [28]. The resulting segmentation covers the neuropil throughout the CNS but does not extend into the surrounding soma regions.

#### Proofreading and reference neurons

Complete neurons were selectively proofread from the whole-CNS FFN segmentation using the Connectome Annotation Versioning Engine (CAVE) [21]. At the time of this evaluation, 24 neurons from the ventral nerve cord (VNC) had been completely proofread. These proofread neurons were used as reference reconstructions for evaluating long-range neuronal continuity.

#### Evaluation

We first compare the three segmentations against the 356 automatically curated Octo reference crops using Variation of Information (VOI) and Adapted Rand Error (ARE) [32, 33]. VOI-split increases when a reference neuron is divided among multiple predicted instances, whereas VOI-merge increases when multiple reference neurons are joined into one predicted instance. Total VOI is the sum of these two terms. ARE measures agreement between predicted and reference instance assignments. Lower values indicate closer agreement for all four measures.

Because these reference crops were derived from the FFN segmentation and then automatically curated, this comparison measures agreement with the curated reference labels rather than performance on an independently annotated test set. As shown in Table 1, FFN has the lowest VOI-split, VOI-merge, total VOI, and ARE across the 356 reference crops. The LSD-based segmentation shows higher split error than the other two methods, while the BiaPy-based segmentation shows higher merge error. Figure 2 provides qualitative comparisons of the three segmentations with the corresponding Octo reference labels.

**Table 1:** Neuron segmentation evaluation. **(a)** Agreement with 356 automatically curated Octo reference crops. Values report mean ± standard deviation across crops. Lower VOI-split, VOI-merge, total VOI, and ARE indicate closer agreement with the (pseudo-)reference labels. **(b)** Long-range continuity evaluated on 24 completely proofread VNC neurons. Higher ERL and nERL indicate greater neuronal continuity.

| (a) Reference-label agreement |  |  |  |  | (b) Whole-neuron continuity |  |
| --- | --- | --- | --- | --- | --- | --- |
| Method | VOI-split ↓ | VOI-merge ↓ | VOI ↓ | ARE ↓ | ERL ($\mu\text{m}$) ↑ | nERL ↑ |
| FFN-based | **0.055 ± 0.032** | **0.008 ± 0.009** | **0.064 ± 0.035** | **0.008 ± 0.010** | **25.287** | **0.0503** |
| LSD-based | 0.514 ± 0.223 | 0.079 ± 0.153 | 0.594 ± 0.300 | 0.065 ± 0.079 | 9.739 | 0.0194 |
| BiaPy-based | 0.139 ± 0.081 | 0.315 ± 0.387 | 0.454 ± 0.391 | 0.127 ± 0.151 | 11.056 | 0.0220 |

**Fig. 2:**
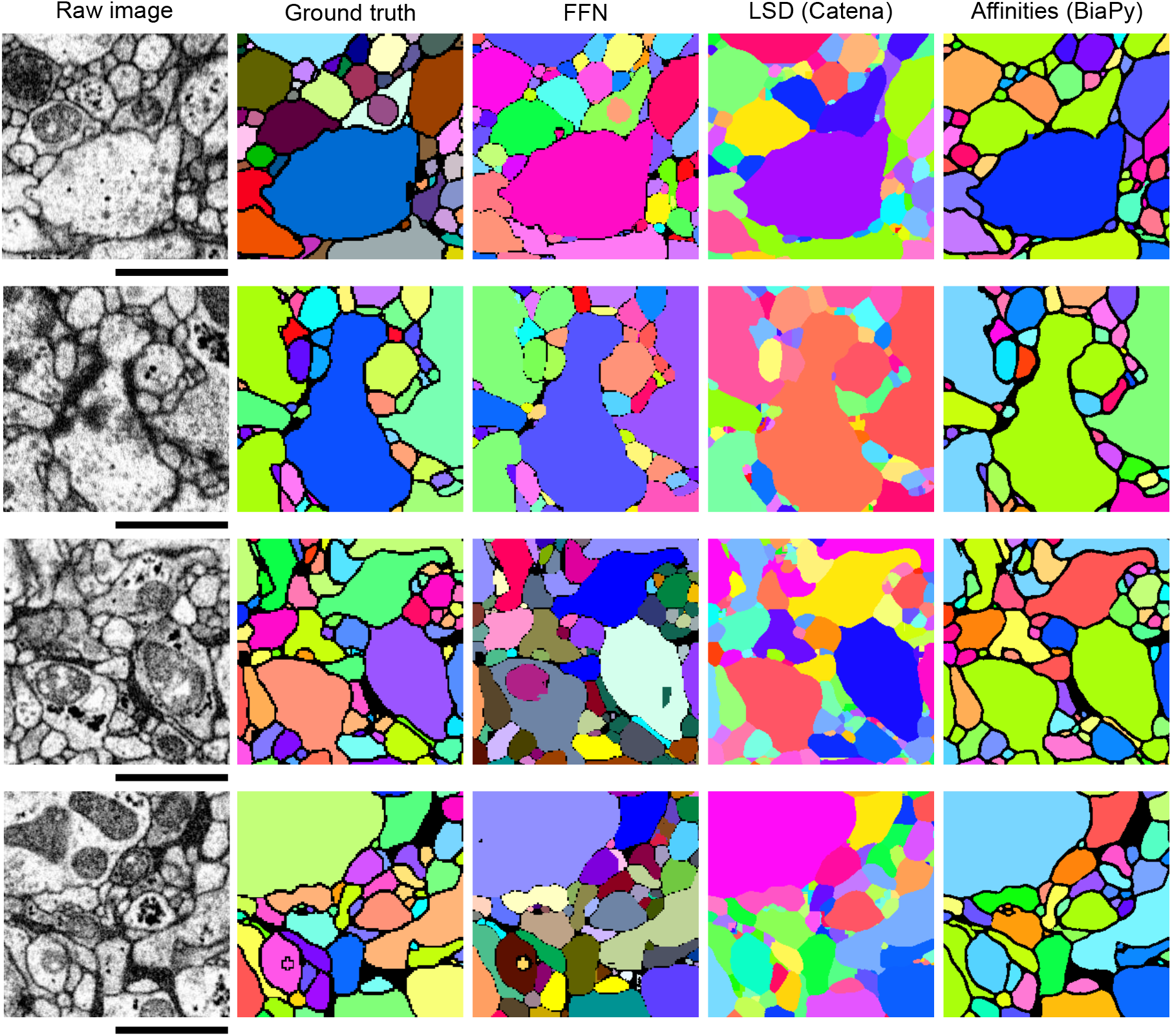
Qualitative comparison of neuron segmentations. Representative Octo neuropil regions showing the raw EM, reference labels, and corresponding FFN-, LSD-, and BiaPy-based neuron segmentations. Colors denote distinct neuron instances. Scale bar: 1 *µ*m.

We additionally evaluate long-range continuity using 24 completely proofread VNC neurons. Expected run length (ERL) [24] measures the expected neurite length that can be followed before a split or merge error interrupts the reconstruction. Normalized ERL (nERL) divides ERL by the maximum achievable ERL for the same set of reference neurons. FFN achieves the highest ERL (25.287 *µ*m) and nERL (0.0503), followed by the BiaPy-based segmentation with 11.056 *µ*m ERL and the LSD-based segmentation with 9.739 *µ*m ERL.

### 3.2 Synaptic Connections

Synaptic connections in Octo were predicted using Synful [34]. Synful predicts directed pre–postsynaptic pairs from a voxel-wise postsynaptic mask and a three-dimensional direction field pointing from postsynaptic locations toward their corresponding presynaptic sites. The original Synful pipeline was refactored and reimplemented in Catena, and we use this implementation for synapse prediction in Octo [27].^4^

#### Synapse annotation and refinement

Synaptic pairs were manually annotated in three 512^3^-voxel Octo subvolumes sampled from different brain regions. The initial annotations contained 542 pairs in Cube 1,870 in Cube 2, and 850 in Cube 3. A Synful model was trained on these annotations and predictions on the same cubes were used to identify locations for targeted re-examination. This process revealed additional synaptic pairs that were absent from the initial annotations, increasing the curated pair counts to 713, 996, and 1015 in Cubes 1–3, respectively.

Before this refinement, we measured pairwise agreement between Synful predictions and the initial annotations using the evaluation procedure of Buhmann et al. [34]. Predicted and annotated synapses were matched as complete pre–postsynaptic pairs using one-to-one assignment, from which precision, recall, and F1 were calculated. Table 2 reports these values. Because the model was trained and evaluated on the same three cubes, these measurements quantify agreement with the initial annotations during label refinement and are not a held-out estimate of model generalization.

**Table 2:** Synful-guided refinement of Octo synapse annotations. Precision, recall, and F1 measure pairwise agreement between Synful predictions and the initial manual annotations on the three training cubes. The final column reports the number of synaptic pairs after targeted re-examination of the annotations. These measurements are in-distribution and are not held-out performance estimates.

| Region | Initial pairs | Precision | Recall | F1 | Refined pairs |
| --- | --- | --- | --- | --- | --- |
| Cube 1 | 542 | 0.660 | 0.777 | 0.714 | 713 |
| Cube 2 | 870 | 0.722 | 0.853 | 0.782 | 996 |
| Cube 3 | 850 | 0.789 | 0.887 | 0.835 | 1015 |

The refinement increased the number of annotated pairs in all three regions, with the largest absolute increase in Cube 1. To assess synaptic partner detection after refinement, a new Synful model was trained on the refined annotations from Cubes 2 and 3 and evaluated on the refined Cube 1. The model achieves a precision of 0.599, recall of 0.998, and pairwise F1 of 0.749 (Table 3).

**Table 3:** Synaptic partner detection after annotation refinement. Synful is trained on the refined annotations from Cubes 2 and 3 and evaluated on the refined Cube 1. Precision, recall, and F1 are computed on matched pre–postsynaptic pairs.

| Test region | Precision | Recall | F1 |
| --- | --- | --- | --- |
| Cube 1 | 0.599 | 0.998 | 0.749 |

After completing annotation refinement and evaluation, a final Synful model was trained on the combined refined annotations from all three cubes and used for whole-CNS inference.

#### Whole-CNS synapse prediction

The final model was applied across the complete Octo CNS to generate predicted pre–postsynaptic pairs. To associate these predictions with the neuron segmentation, the FFN segment ID was queried at each predicted pre- and postsynaptic site, and the two sites were associated with the corresponding neuronal segments. Predictions falling on background were discarded, as were pairs for which both sites were assigned to the same FFN segment. Because the FFN segmentation has not been exhaustively proofread, these segment IDs primarily correspond to neuronal fragments rather than complete neurons.

After this filtering and neuron assignment, the Octo reconstruction contains approximately 3.5 million putative synaptic pairs. These predictions form the synaptic layer released with Octo. Figure 3 shows predicted synaptic sites along an example reconstructed neuron together with EM slices at three selected locations.

**Fig. 3:**
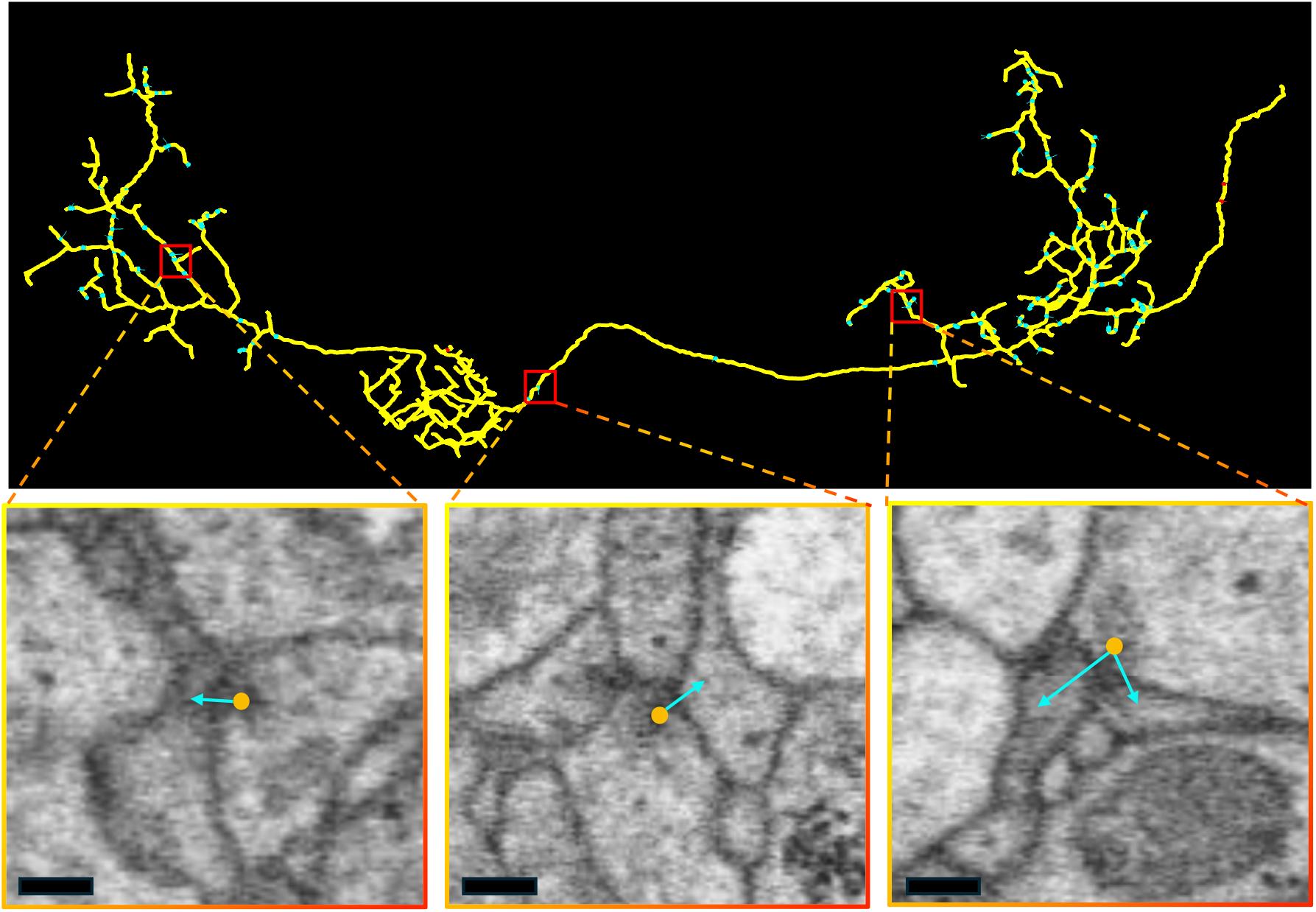
Predicted synaptic connections in Octo. Top, an example reconstructed neuron with predicted synaptic sites, with three locations highlighted. Bottom, corresponding EM slices of the highlighted synapses showing the presynaptic site (orange ball) and its corresponding postsynaptic partner (cyan arrow). Scale bar: 500 nm.

### 3.3 Mitochondria Segmentation

We also release a whole-CNS mitochondria instance segmentation containing 1,262,478 predicted instances. The segmentation was generated using a residual U-Net (RSU-Net) adapted from the architecture of Xie et al. [35] for isotropic FIB-SEM data. In particular, the anisotropy-aware pooling and convolution strides of the original architecture were replaced with isotropic operations to match the 8 × 8 × 8-nm Octo resolution. The network predicts three voxel-wise representations: a mitochondria foreground mask, a signed distance-based representation, and a contour map. The distance and contour predictions act as auxiliary representations that improve foreground delineation, while the final mitochondria instances are obtained from the predicted foreground mask using connected-component labeling.

#### Training and evaluation

Our RSU-Net model was trained using five 512^3^-voxel Hemibrain volumes together with an annotated Octo volume of shape 300 × 1269 × 1572 voxels in *z, y, x*. Approximately 10% of the Octo training volume was reserved for validation, while a separate Octo volume of shape 200 × 1269 × 1572 voxels was held out for evaluation. CutMix augmentation [36] was used during training, with checkpoint selection and early stopping based on the Octo validation data.

We evaluate both foreground recovery and instance correspondence on the held-out Octo volume. IoU and Dice measure voxel-wise foreground overlap, while the association metrics characterize one-to-one correspondences, over-segmentation, under-segmentation, many-to-many associations, and missed instances following Franco-Barranco et al. [37]. As shown in Table 4, the selected RSU-Net achieves an IoU of 0.7431 and Dice of 0.8526. At the instance level, 69.91% of reference mitochondria have a one-to-one association with a prediction. The largest remaining error category is under-segmentation (17.02%), corresponding to cases where multiple reference mitochondria are merged into a single predicted instance.

**Table 4:** Mitochondria segmentation on held-out Octo annotations. Semantic metrics measure foreground overlap, while association metrics describe the correspondence between predicted and reference mitochondria instances.

| Metric | Value |
| --- | --- |
| *Semantic overlap* |  |
| IoU | 0.7431 |
| Dice | 0.8526 |
| *Instance associations* |  |
| One-to-one | 69.91% |
| Over-segmentation | 5.17% |
| Under-segmentation | 17.02% |
| Many-to-many | 1.22% |
| Missed instances | 6.69% |

Figure 4 shows representative EM views together with the corresponding predicted mitochondria instances from the released whole-CNS segmentation.

**Fig. 4:**
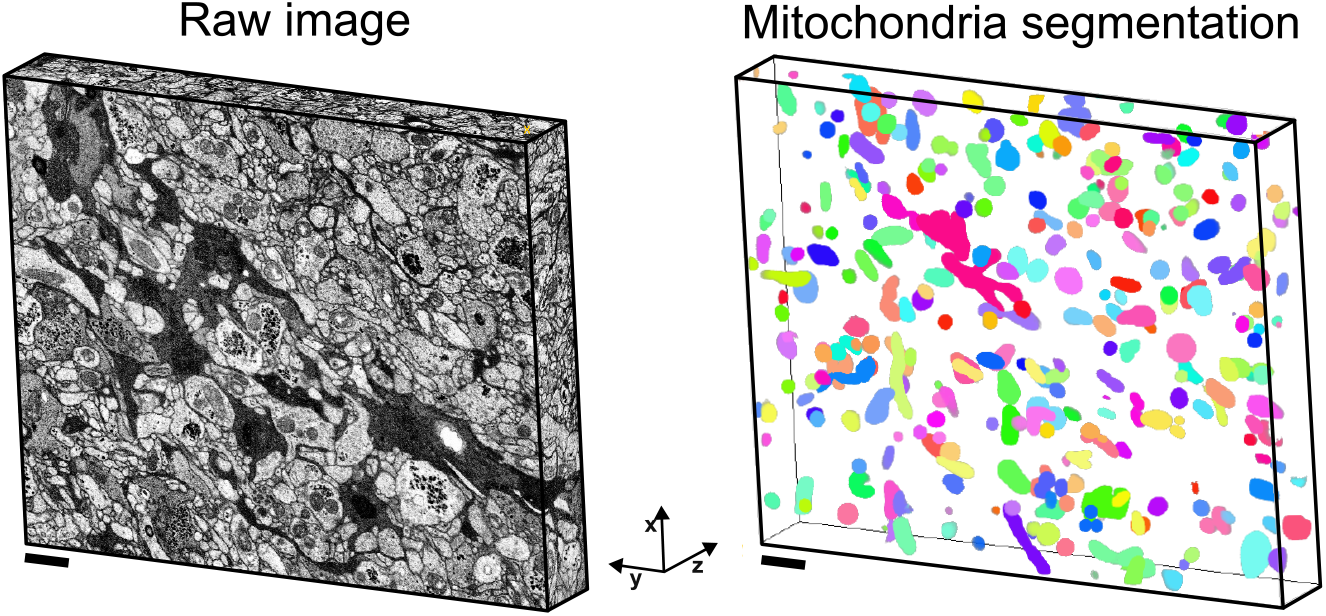
Mitochondria segmentation in Octo. Representative EM views and corresponding predicted mitochondria instances from the released whole-CNS segmentation. Scale bar: 1 *µ*m.

### 3.4 Glia Segmentation

Glial processes closely interact with neuronal processes throughout the CNS. In the larval brain, astrocyte-like glia extend processes into synaptic neuropil, whereas ensheathing glia organize around and surround neuropil regions. To capture this complementary anatomical context, we also release a whole-CNS semantic segmentation of glial tissue in Octo, alongside the neuron reconstructions and predicted synaptic sites.

#### Training and evaluation

Ground-truth glia masks were manually curated in an Octo subvolume of shape 502*×*1269*×*1572 voxels in *z, y, x*. This volume was divided into a training region of shape 300*×*1269*×*1572 and a held-out test region of shape 202*×*1269*×*1572, with 10% of the training data reserved for validation.

We extend the same three-dimensional residual U-Net (RSU-Net) architecture used for mitochondria segmentation (Sec. 3.3) to semantic glia segmentation. The model was trained on the glia annotations using CutMix augmentation [36], with model selection and early stopping based on the validation split.

On the held-out Octo region, the model achieves a Dice score of 0.8489 and an IoU of 0.7375, as reported in Table 5. The selected model was subsequently applied across the complete Octo CNS to generate the released semantic glia segmentation.

**Table 5:** Glia segmentation on held-out Octo annotations. Semantic segmentation performance on the 202 *×* 1269 *×* 1572-voxel held-out test region.

| Metric | Value |
| --- | --- |
| Dice | 0.8489 |
| IoU | 0.7375 |

Figure 5 shows a representative EM section together with the corresponding predicted glia mask from the whole-CNS segmentation.

**Fig. 5:**
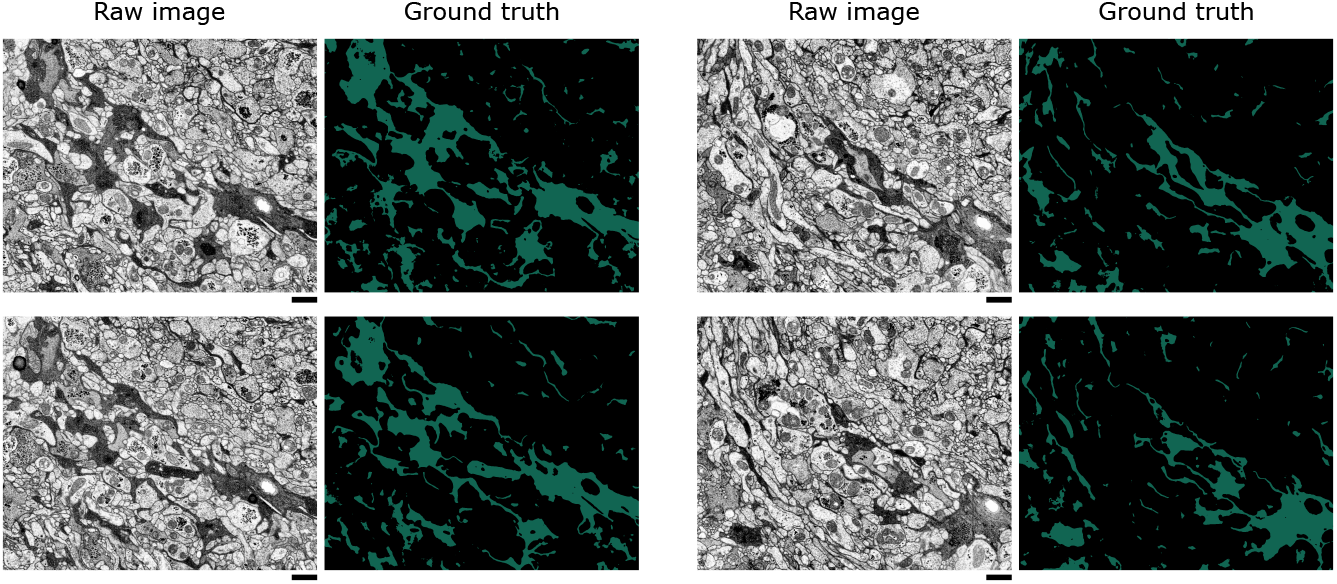
Glia segmentation in Octo. Representative raw EM section containing glial tissue and the corresponding predicted semantic glia mask from the whole-CNS segmentation. Scale bar: 1 *µ*m.

## 4 Data Availability

Evaluation datasets associated with this work are publicly available on Zenodo at https://zenodo.org/records/22832866.

The Octo EM volume and released neuron segmentation layers can be explored interactively in Neuroglancer at https://tinyurl.com/octo-seg.

The released mitochondria and glia segmentations are available at https://tinyurl.com/mito-glia.

The predicted synaptic connections are available through the CAVE instance associated with the Octo reconstruction. Access to CAVE and CATMAID for proofreading, annotation, and inspection of the synaptic data is available upon request. Please contact the authors for access.

## 5 Code Availability

Code used in this work is publicly available through Catena at https://github.com/Mohinta2892/catena and BiaPy at https://github.com/BiaPyX/BiaPy.

## 7 Acknowledgments

This project was funded by the HHMI Janelia Research Campus, a Wellcome Trust Investigator Award to A.C. (Ref: 205038/Z/16/Z), an ERC grant to M.Z. (Ref: ERC-2018-COG: 819650), a BBSRC Grant to A.C. (Ref: APP26929, “Population connectomics”) and MRC LMB core funding to M.Z. and A.C.

## Footnotes

1 Registration scripts: https://github.com/acardona/scripts/blob/e8a34edde0a02d905503c2f8ccbbaa88999ec969/python/imagej/FIBSEM/serial_section_registration.py.

2 FFN model training and inference on the Octo volume were performed by the Google Connectomics team.

3 https://github.com/Mohinta2892/catena

4 https://github.com/Mohinta2892/catena

